# Role of RIPK1 in Diffuse Gliomas pathology

**DOI:** 10.1101/2023.11.11.566709

**Authors:** Leslie C. Amorós Morales, Santiago M. Gómez Bergna, Abril Marchesini, Ma. Luján Scalise, Nazareno González, Marianela Candolfi, Víctor Romanowski, Matias L. Pidre

**Affiliations:** Instituto de Biotecnología y Biología Molecular (IBBM-CONICET-UNLP), Facultad de Ciencias Exactas, Universidad Nacional de La Plata, La Plata, Argentina; Instituto de Investigaciones Biomédicas (INBIOMED-CONICET-UBA), Facultad de Ciencias Médicas, Universidad de Buenos Aires, Buenos Aires, Argentina

**Keywords:** Glioma, RIPK1, Pyroptosis, EMT, tumor-infiltrating immune cells

## Abstract

**Purpose:** The aim of the present work was to investigate the role of Receptor-interacting protein kinase 1 (RIPK1) both in mutated and wild type isocitrate dehydrogenase (IDH) Diffuse Gliomas (DG).

**Patients and Methods:** We analyzed RIPK1 mRNA expression in DG databases from The Cancer Genome Atlas (TCGA) containing clinical, genomic and transcriptomic information from 661 patients. Transcriptomic studies (mRNA expression levels, correlation heatmaps, survival plots and Gene Ontology and meta-analysis of immune gene signatures) were performed with USC Xena and R. Statistical significance was set at p-values less than 0.05.

**Results:** The results showed a lower survival probability in patients belonging to the high RIPK1 expression subgroup compared to those samples with low RIPK1 expression. We also observed a higher expression of RIPK1 in wtIDH samples compared to those with mIDH. In order to further characterize the role of RIPK1 in DG, we performed a Gene Ontology and Pathway Enrichment Analysis using the Xena platform’s differential expression tool. The results showed that RIPK1 is involved in inflammatory and immune responses. Hence, the expression levels of some of the genes involved in the following molecular processes crucial for cancer progression were studied: proliferation, epithelial-mesenchymal transition, immune cell infiltration and cell death pathways. Briefly, the results showed significant differences in genes related to increased cellular dedifferentiation, proinflammatory cell death pathways and tumor infiltrating immune cells gene signatures (Welch’s t-test).

**Conclusion:** RIPK1 over-expression is associated with a poor prognosis in DG. This fact, together with our results suggest that RIPK1 may play a crucial role in glioma pathogenesis highlighting the need to take into account RIPK1 expression levels for decision making when choosing or designing therapeutic alternatives.

**CONTEXT SUMMARY:** *Key Objective:* Evaluate the role of the Receptor-interacting protein kinase 1 (RIPK1) in Diffuse Gliomas (DG) pathology through an exhaustive *in silico* patient database analysis.

*Knowledge generated:* We demonstrated that RIPK1 is overexpressed in more aggressive DG and correlates with clinical attributes associated with poor prognosis. In addition, our analyses showed that high RIPK1 expression correlates with key genes involved in pro inflammatory cell death pathways and an increased expression of immune gene signatures suggesting greater immunological infiltration in the tumor.

*Relevance:* Our results from patient database analyses propose RIPK1 as a new relevant molecular prognosis marker for DG. Our findings are in concordance with different preclinical studies and provide additional information that can be useful for decision making when choosing therapeutic strategies and for the development of novel therapeutic approaches such as gene or immunotherapy.

*This work was presented in:* XIII Argentine Congress of Bioinformatics and Computational Biology (XIII CAB2C), XIII International Conference of the Iberoamerican Society of Bioinformatics (XIII SoIBio) and III Annual Meeting of the Ibero-American Artificial Intelligence Network for Big BioData (III RiaBio).

## BACKGROUND

Diffuse gliomas (DG) are the most frequent malignant primary brain tumors in adults, constituting 50% of them ^1, 2^. These tumors originate from glial cells and are highly infiltrative and heterogeneous.

Despite advances in conventional therapies (neurosurgery, radiotherapy, and chemotherapy), most patients with DG eventually die due to its highly invasive nature and intrinsic resistance to therapies. For this reason, it is imperative to identify new molecular targets to develop new treatment strategies ^3^.

The classification of DG has traditionally been based on their histopathologic features ^4^. However, in 2016 the WHO included for the first time distinctive genetic and epigenetic alterations to define various groups of gliomas ^5–8^. Among all characterized genetic alterations, the mutational status of isocitrate dehydrogenase (IDH) 1 and 2 has become the main marker for the molecular classification and prognosis of DG in adults. IDH, an enzyme involved in cell metabolism and the response to oxidative stress ^9^, catalyzes the conversion of isocitrate to α-ketoglutarato (αKG). A mutation in the R132 residue of the active site modifies the catalytic activity of the enzyme, which converts αKG to 2-hydroxyglutarate (2-HG) ^10–12^. The accumulation of 2-HG leads to hypermethylation-associated epigenetic reprogramming of tumor cells ^11, 12^. Although the IDH mutation (mIDH) is associated with a poor prognosis in some cancers such as leukemia ^13^, the opposite has been observed in DG ^14, 15^. The IDH1 and IDH2 mutation is a positive prognostic marker in patients with glioma and is associated with a higher survival rate when compared with wild-type IDH (wtIDH) patients ^3, 14, 15^.

Apoptosis, necroptosis, and pyroptosis play a key role in tumor homeostasis ^16^. Since Apoptosis has been widely studied as an important anticancer defense mechanism, the role of necroptosis and pyroptosis in cancer is not fully understood at present. Pyroptosis is closely related to nervous system diseases, infectious diseases, autoimmune diseases, cardiovascular diseases, and tumors. With further research, the relationship between pyroptosis and tumors is becoming increasingly clear, which provides some strategies for clinical treatments ^17^.

Receptor-interacting serine/threonine protein kinase 1 (RIPK1) is a master regulator of the cellular decision between pro-survival NF-κB signaling and death in response to a broad set of inflammatory stimuli ^18–20^.

Necroptosis is primarily mediated by RIPK1, RIPK3, and MLKL (mixed lineage domain-like kinase pseudokinase) and inhibited by necrostatin-1 (Nec-1) ^21^.

In addition to its key role in viral infection, necroptosis has been suggested to play a critical role in the regulation of cancer biology, including cancer oncogenesis, metastasis, and immunity ^22, 23^. Dual effects of necroptosis on cancer have been observed. On the one hand, it has been suggested that key mediators of the necroptotic pathway alone or in combination promote metastasis and tumor progression ^24–26^. However, necroptosis also serves as an alternative mechanism that protects against tumor development when apoptosis is compromised ^27, 28^. Given the ambiguous role of necroptosis in cancer biology, necroptosis has emerged as a novel target for cancer therapy, and an increasing number of compounds inducing or manipulating necroptosis are in the pipeline of development ^29, 30^.

In this work we set out to characterize the role of RIPK1 in the tumor progression and the tumor immune microenvironment (TIME) of diffuse gliomas through exhaustive *in silico* analyzes from patient databases.

## METHODS

### Glioma patient samples

For the analysis we used public datasets containing clinical, genomic and transcriptomic information from patient samples. We also used two different platforms to analyze the data. For all the analyses, the TCGA-LGGGBM database and the UCSCXena platform were employed. Samples corresponding to DG were filtered, classified according to IDH status and the expression of RIPK1 and other genes was evaluated.

### RIPK1 expression and clinical attributes

For these analyses, the glioma database TCGA-LGGGBM and the UCSCXena^31^ platform were used. RIPK1 was used as a query and the samples were divided into two groups according to the median value of expression: high expression of RIPK1 (>9.094) and low expression of RIPK1 (≤9.094). Correlation between RIPK1 expression levels and several clinical attributes of interest was assessed.

### Survival plots

The UCSCXena^31^ platform and the TCGA-LGGGBM database were used to analyze the patient’s survival. For this, samples corresponding to different types of glioma were filtered and the expression of RIPK1 were evaluated. In the analysis, the samples were divided into two groups according to the median value of expression: high expression of RIPK1 (>9.094) and low expression of RIPK1 (≤9.094), and survival was plotted for each group.

### Transcriptomic analysis

The UCSCXena^31^ and cBioPortal^32, 33^ platforms and the TCGA-LGGGBM database were used to evaluate the expression of RIPK1 ^34^. The mRNA expression data in z-Scores relative to all samples (log microarray) carried out on the Illumina HT-12 v3 platform (Illumina_Human_WG-v3) was used.

For different pathway correlation analyses, the TCGA and the UCSCXena and cBioPortal platforms were used.

### Gene Ontology and pathway analysis

TCGA-LGGGBM dataset was used to perform differential gene expression analysis and Gene Ontology. The dataset contained fully sequenced transcriptome data from 702 primary tumor samples from glioma patients.

To perform the differential expression analysis of genes, the samples were separated into two groups: RIPK1 expression values greater than the median and RIPK1 expression values lower or equal than the median. Differential expression analysis with subsequent Gene Ontology and pathway enrichment analysis was performed using the extension available in the Xena platform with default parameters comparing both groups.

### Meta-analysis of immune gene signatures in mIDH and wtIDH glioma biopsies

A total of 521 patients with gliomas from the TCGA-LGGGBM cohort were analyzed in this study. We stratified patients according to the mutational status of IDH into two groups: mIDH and wtIDH patients. Gene signatures for each immune cell type were obtained from previously published data ^35, 36^. Samples were classified into two groups according to RIPK1 mRNA expression levels in RIPK1low and RIPK1high using median expression values as cut-off. Normalized expression values were log2-transformed. Genes were grouped into 8 immune cell type signatures (Cytotoxic CD8, Helper T cells, Dendritic cells, NK, Macrophages, Neutrophils, Memory T cells and Treg), derived from Gonzalez et al.^3^ ESTIMATE scores were obtained from the cBio Cancer Genomics Portal ^32, 33^. Gene signatures for each immune cell type were obtained from previously published data ^37, 38^.

### Statistical analysis

Expression of RIPK1 in samples of TCGA-LGGGBM vs. different DG subtypes: n = 1106; Kruskal-Wallis test followed by the Dunn test with the P value corrected by the Bonferroni method. Expression of RIPK1 in samples of TCGA_LGG_GBM vs. radiotherapy treatment: n = 701; Welch’s t-test. Survival plots: n = 702; logRank Test. RIPK1 expression vs. proteins of different pathways: n = 702 (n = 351 High Expression of RIPK1; n = 351 Low Expression of RIPK1); Welch’s t-test. Correlation analysis: Pearson’s correlation coefficient. Meta-analysis of immune gene signatures in mIDH (n=311) and wtIDH (n=210): Mann-Whitney test. Statistical significance was considered when p-value did not exceed 0.05 for all studies.

## RESULTS

### Patient cohort

For clinical and transcriptomic analysis TCGA-LGGGBM database was consulted. Patient attributes are summarized in Table 1.

**Table 1.** Patient cohort. Table 1 shows principal clinical attributes corresponding to the TCGA LGGGBM database used in this work. The number in each cell indicates the quantity of patients with clinical attributes specified in column one.

| Clinical attribute |  | Database: TCGA<br>LGGGBM |
| --- | --- | --- |
| Total of patients |  | 670 |
| RIPK1 | High expression | 335 |
|  | Low expression | 335 |
| IDH status | wtIDH | 257 |
|  | mIDH | 404 |
|  | Undefined | 9 |
| Age at initial<br>pathologic<br>diagnosis | <50 | 383 |
|  | >50 | 275 |
|  | Undefined | 2 |
| Histological<br>classification | Oligoastrocytoma | 130 |
|  | Oligodendroglioma | 191 |
|  | Astrocytoma | 324 |
|  | GBM | 153 |
| Radiotherapy | Yes | 408 |
|  | No | 200 |
|  | Discrepancy | 1 |
|  | Undefined | 61 |

### RIPK1 expression and survival

The survival of patients with glioma was evaluated on 670 samples from TCGA Pan-Cancer Atlas. First, samples were separated into two groups, high and low RIPK1 expression, and a survival was plotted for both groups. A significantly lower survival probability was observed in patients with higher RIPK1 expression (Figure 1.A).

**Figure 1.**
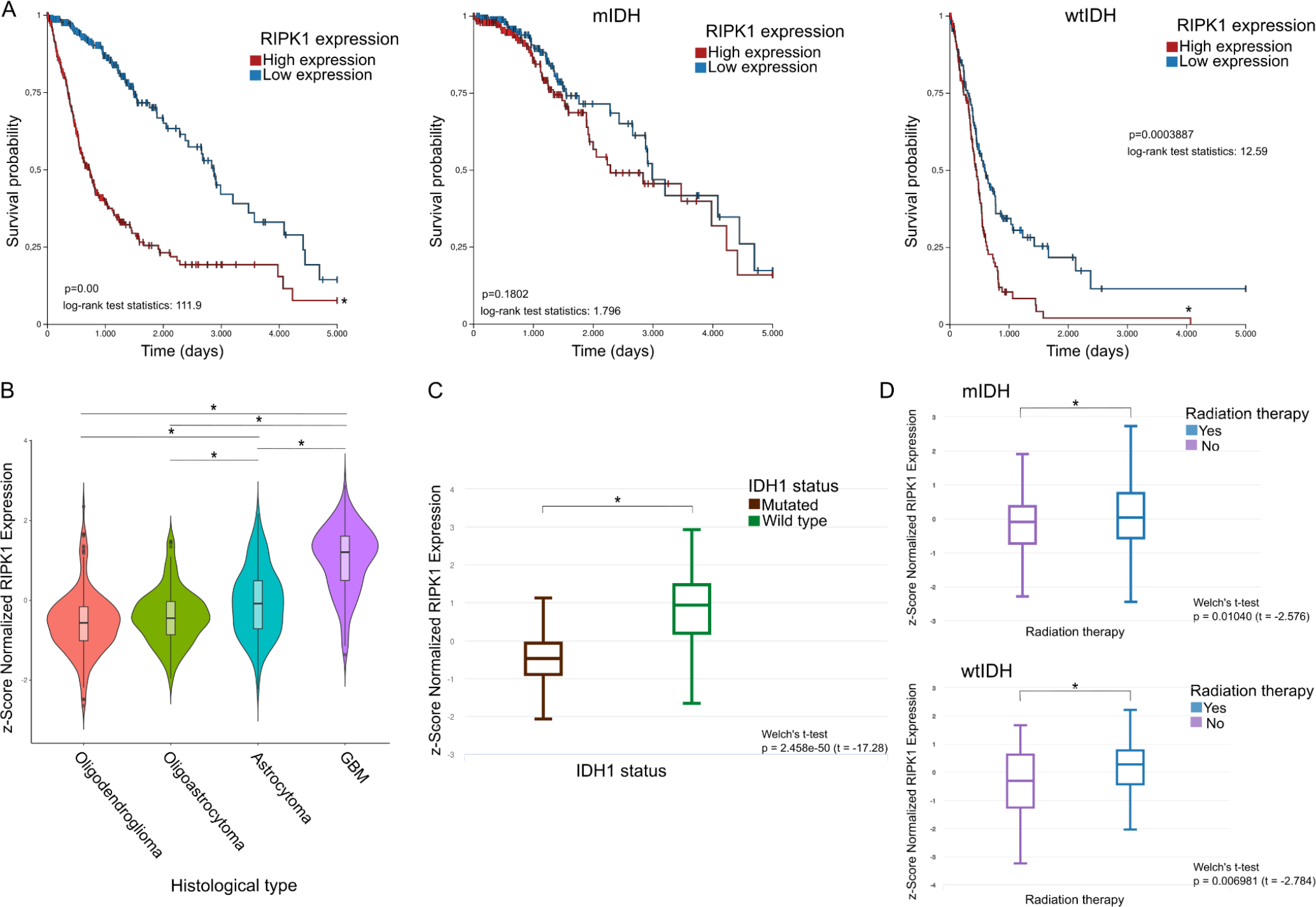
**(A)** Survival plots. DG samples were filtered from TCGA, and the expression of RIPK1 was evaluated. In the analysis, the samples were divided into two groups according to the median value of expression: high expression of RIPK1 and low expression of RIPK1 and survival was plotted for each group. Data were plotted unclassified (left) or classified into mIDH (center) or wtIDH (right). Log-rank test *P < .05. **(B)** Differential expression of RIPK1 between DG tumor subtypes. Kruskal-Wallis test followed by the Dunn test with the P value corrected by the Bonferroni method. *P < .05. **(C)** Differential expression of RIPK1 in mIDH and wtIDH gliomas. Welch’s T-test. *P < .05. **(D)** Differential expression of RIPK1 between samples of patients who received radiotherapy and those who did not. Samples were classified into mIDH (top) and wtIDH (bottom). Welch’s T-test. *P < .05.

For a more exhaustive analysis, samples were filtered by IDH status, mutated or wild type, before being divided based on RIPK1 expression. Hereby, 404 samples from patients with mutated IDH1 were analyzed and no significant differences in survival probability were observed under conditions of either low and high expression of RIPK1 (Figure 1.A). However, in those samples with wild type IDH1, the survival probability was decreased in the high RIPK1 expression group (Figure 1.A).

RIPK1 expression was evaluated in samples derived from patients with different subtypes of glioma using cBioPortal platform. The results showed higher levels of RIPK1 in samples derived from patients with GBM and Astrocytoma CNS grade 2–3 (A2-3) compared with samples from low grade glioma, such as oligodendroglioma (OD) and wtIDH gliomas grade 2–3 (Figure 1.B).

To evaluate RIPK1 expression and IDH status, samples were divided by wild type or mutated IDH. Thus, a significant increase in the expression of RIPK1 in wtIDH in comparison with mIDH was observed (Figure 1.C).

On the other hand, samples were divided according to patients who had received radiotherapy and those who had not and RIPK1 expression was studied in both cases. A higher expression of RIPK1 was observed in samples derived from patients treated with radiotherapy compared with samples from untreated patients (Figure 1.D).

### RIPK1 and pathways

After classifying the samples from the TCGA-LGGGBM database according to the median expression of RIPK1, Differential Expression Analysis was performed followed by Gene Ontology (Figure 2). This first analysis showed that in those samples with high expression of RIPK1, the pathways linked to cell dedifferentiation, inflammation and cell death are upregulated.

**Figure 2.**
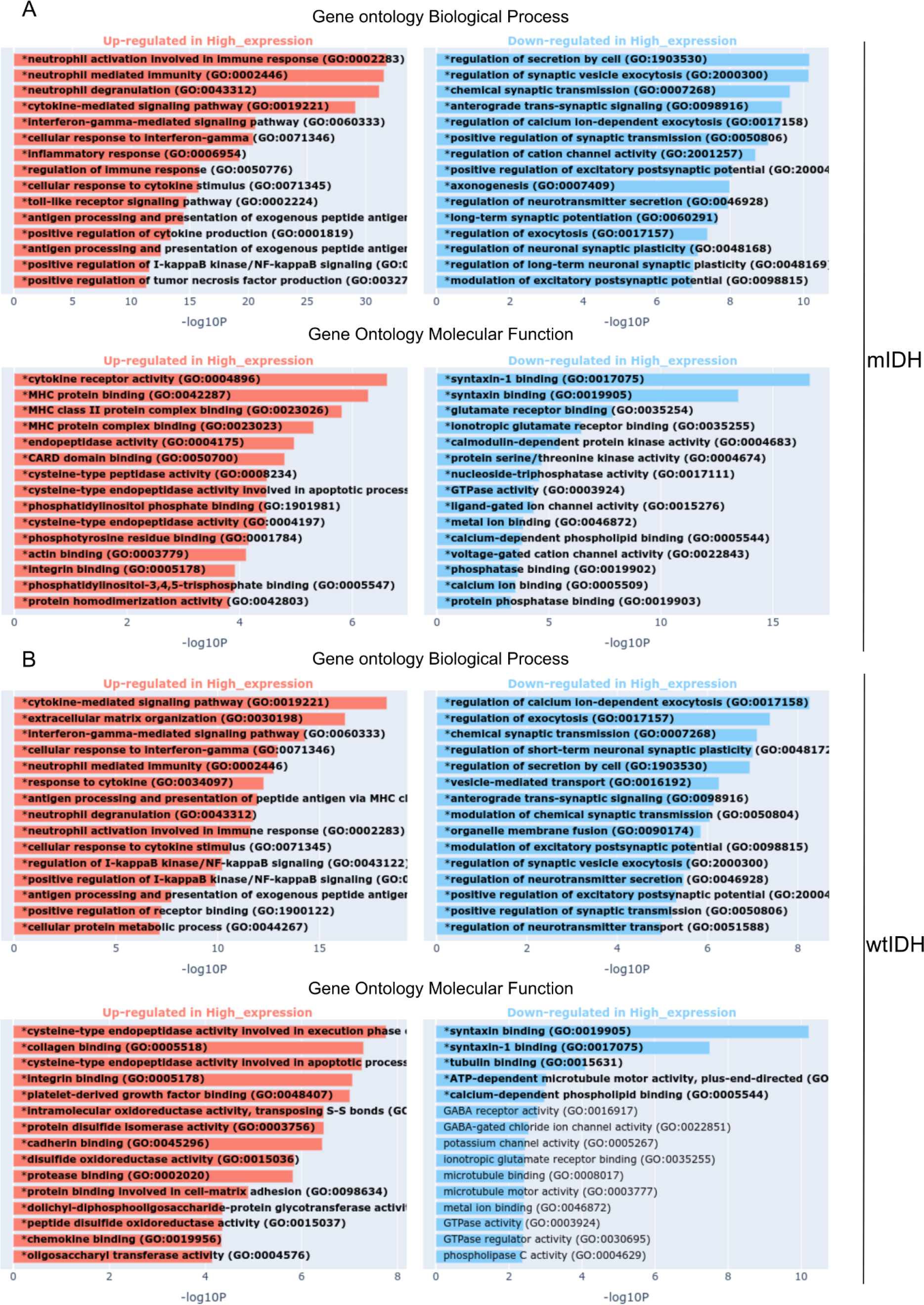
Gene ontology and over-represented pathways. Graphs show the canonical pathways significantly over-represented (enriched) by the DEGs with the number of genes for the main categories of the three ontologies (GO: biological process, GO: cellular component, and GO: molecular function, respectively). DEA genes low expression versus RIPK1 high expression: **(A)** mIDH samples and **(B)** wtIDH samples. The statistically significant canonical pathways in the DEGs are listed according to their P value corrected by FDR (–log10; colored bars) and the ratio of the listed genes found in each pathway over the total number of genes in that pathway. The information was obtained from The Cancer Genome Atlas Pan-Cancer-Xena database. DEA, differential expression analysis; DEG, differential expressed gene; FDR, false discovery rate.

### RIPK1 and cellular homeostasis

#### i. Apoptosis

Differential expressions of apoptosis related genes were analyzed in both groups, high and low RIPK1 expression. For a further analysis, samples were divided, or not, by IDH1 status and correlation between RIPK1 expression and apoptosis related genes was studied. It was observed that BAX, CASP3, CASP8, TP53, CYCS and NFкB1 have significantly higher levels in the group with high RIPK1 expression compared with those with lower levels of RIPK1. On the other hand, the results showed a lower expression of CASP9 in the high RIPK1 expression group while no significant differences were observed for BCL2 and DIABLO (Figure 3.A).

**Figure 3.**
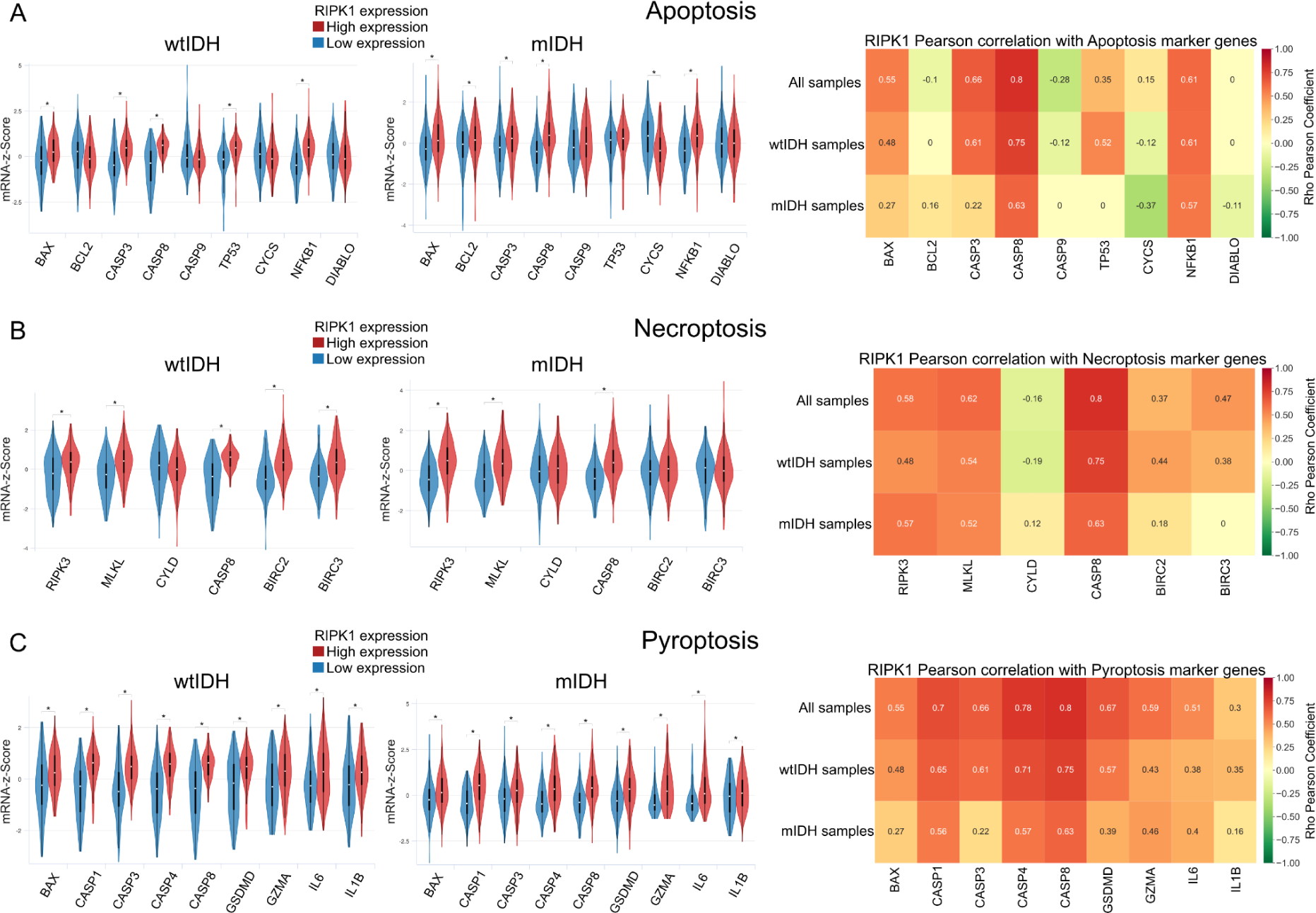
Transcriptomic analysis (TCGA) of **(A)** Proliferation and **(B)** Epithelial-mesenchymal transition (EMT). In each panel, Left: Boxplot of the expression (z-score) of proteins involved in the respective pathway under conditions of high and low RIPK1 expression. Samples were classified in mIDH and wtIDH and then plotted. Welch’s t-test *P <.01. Right: Correlation between RIPK1 expression and proteins involved in the respective pathway separated in all samples, mIDH and wtIDH tumors. Numbers in cells indicate the correlation Pearson index in statistically significant comparisons.

#### ii. Necroptosis and Pyroptosis

Genes related to necroptosis, a regulated form of necrosis, were analyzed under conditions of either low or high RIPK1 expression. The results showed higher levels of RIPK3, MLKL, CYLD, CASP8 and BIRC3 in those samples with high RIPK1 expression while no significant differences were observed for CYLD. Also, the correlation between necroptosis related genes and RIPK1 expression was studied in three different groups: samples with mutated IDH1, those samples with wild type IDH1 and a group that contained both status (Figure 3. B)

A similar analysis was performed to study pyroptosis, a proinflammatory regulated cell death. In this case, BAX, CASP1, CASP3, CASP4, CASP8, GSDMD, GZMA, IL6 and IL1β were evaluated and it was observed a higher expression of all the previously mentioned genes in the samples with high RIPK1 expression compared with those with low RIPK1 expression (Figure 3.C).

#### iii. Proliferation

Cell proliferation is a vital process to tumor progression. Thus, transcriptomic analysis was repeated evaluating genes linked to this process under conditions of low and high RIPK1 expression. In addition, the correlation between those genes and RIPK1 expression was studied dividing, or not, the samples by IDH1 status. The results showed a higher expression of MAPK8, MAP2K1, MAPK1, MAPK3, APC, PTPN11, CSNK2A1, GSK3B, AXIN1, AXIN2, KRAS and a lower expression of JUN, STAT3, NFKB1, AKT1, CTNNB1, NRAS and NOTCH2 in the group of low RIPK1 expression, while no significant differences were observed for PIK3CA, HRAS and RAF1 (Figure 4.A).

**Figure 4.**
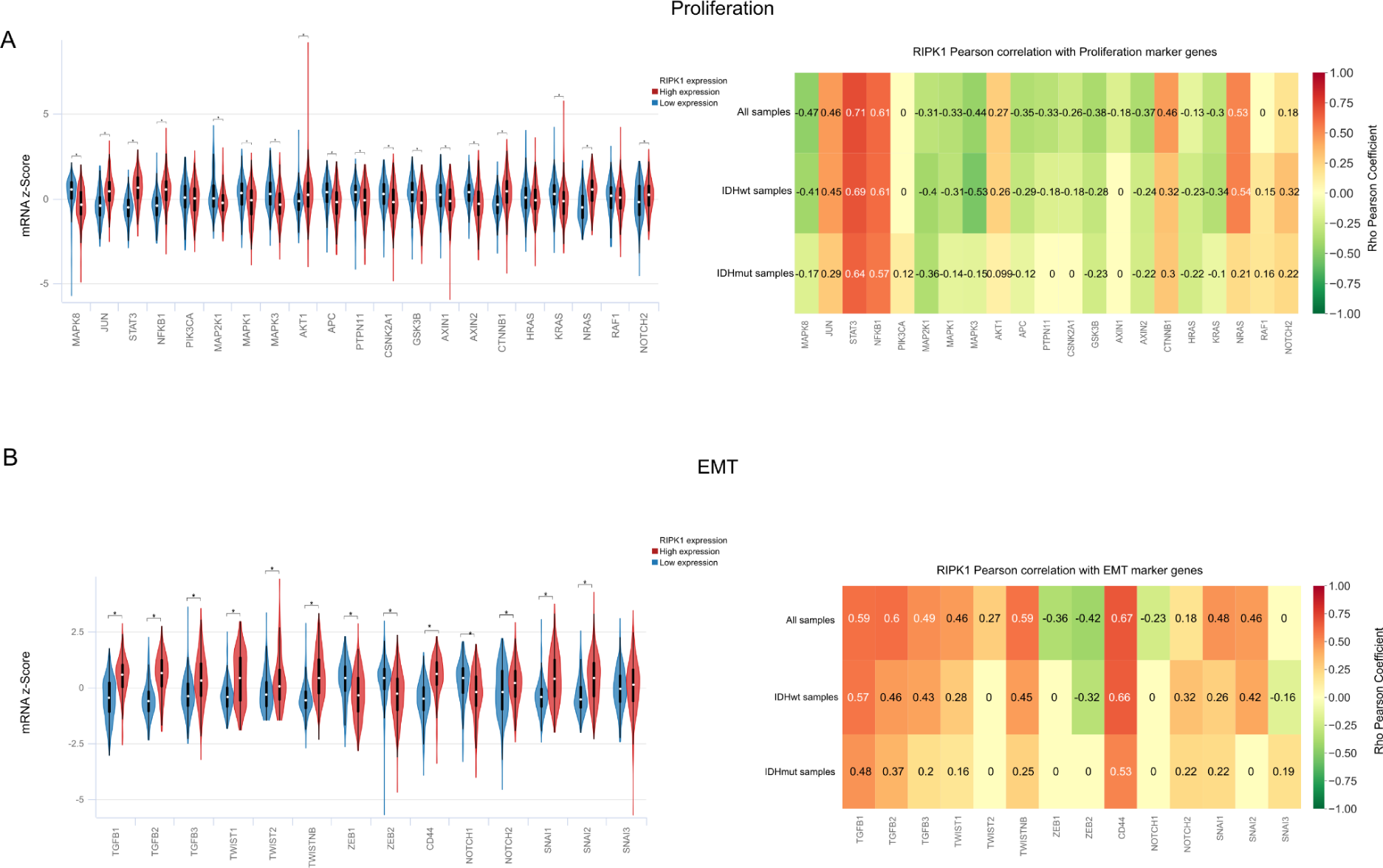
Transcriptomic analysis of cell death pathways (TCGA). **(A)** Apoptosis, **(B)** Necroptosis and **(C)** Pyroptosis. In each panel, Left: Boxplot of the expression (z-score) of proteins involved in the respective pathway under conditions of high and low RIPK1 expression. Samples were classified in mIDH and wtIDH and then plotted. Welch’s t-test *P <.01. Right: Correlation between RIPK1 expression and proteins involved in the respective pathway separated in all samples, mIDH and wtIDH tumors. Numbers in cells indicate the correlation Pearson index in statistically significant comparisons.

### RIPK1 and EMT

In the context of neoplasias, epithelial-mesenchymal transition is associated with tumor initiation, invasion and metastasis ^39^. In this case, TGFB1, TGFB2, TGFB3, TWIST1, TWIST2, TWISTNB, ZEB1, ZEB2, CD44, NOTCH1, NOTCH2, SNAI1, SNAI2 and SNAI3 were studied in groups of either low and high RIPK1 expression. Finally, the study of the correlation between RIPK1 expression and the expression of the previously mentioned genes was carried out in samples with mIDH, wtIDH or both of them. It was observed that samples with high RIPK1 expression presented higher levels of the majority of the evaluated genes, but they presented lower expression of ZEB1, ZEB2 and NOTCH1, while no significant differences were observed for SNAI3 (Figure 4.B).

### RIPK1 expression and immune cell infiltration

Given the potential role of RIPK1 in glioma immunity due to its role in necroptosis and pyroptosis, we performed a meta-analysis using previously validated immune genetic signatures of tumor-infiltrating immune cells ^37, 38^. We found that the expression of genetic signatures of different lymphocyte populations, i.e. helper, cytotoxic, memory and regulatory T cells, was upregulated in mIDH gliomas with high RIPK1 expression levels (Figure 5.A). Similarly, the expression of gene signatures of antigen presenting cells, i.e. dendritic cells (DCs) and macrophages, was upregulated. The same effect is observed regarding neutrophils and NK cells (Figure 5.A).

**Figure 5.**
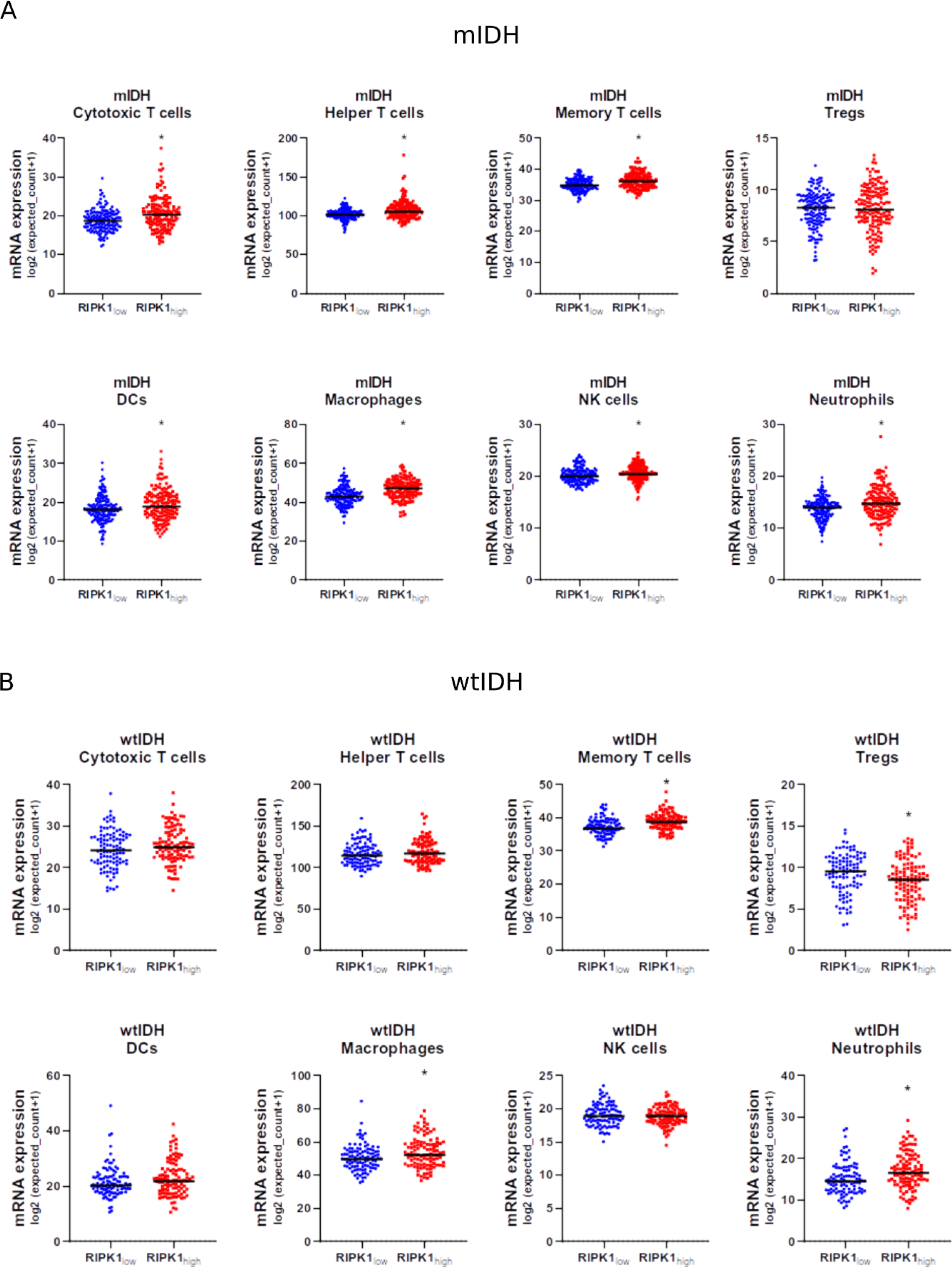
Meta-analysis of immune gene signatures in mIDH and wtIDH glioma biopsies. We evaluated the gene signatures that characterize immune cell populations in transcriptomic data of glioma biopsies deposited in The Cancer Genome Atlas. Data were stratified according to the 2021 WHO classification in **(A)** mIDH (n=311) and **(B)** wtIDH gliomas (n=210). Patients in each group were classified according RIPK1 mRNA expression levels into two groups (RIPK1low and RIPK1high). *, p <0.05; Mann-Whitney test.

In the case of wtIDH gliomas, we observed that the expression of genetic signatures of macrophages and neutrophils was upregulated in high RIPK1 expression levels samples (Figure 5.B). Regarding the expression of genetic signatures of different lymphocyte populations, we only observed an increase in memory T cells. However, we observed a significant decrease in the expression of genetic signatures of Tregs in wtIDH gliomas with high RIPK1 expression levels (Figure 5.B).

## DISCUSSION

Diffuse gliomas are a heterogeneous collection of brain tumors. They are among the most common and deadly types of primary brain tumors ^1, 2^. At the moment, adult gliomas should be divided into two major groups based on the mutational status of the isocitrate dehydrogenase genes IDH1 or IDH2. IDH-mutant gliomas typically present as lower histologic grades with improved prognosis ^40^, but they often evolve to higher grades. In contrast, IDH–wild-type gliomas usually present as glioblastomas (GBM), the most common and clinically aggressive World Health Organization (WHO) grade IV gliomas, with median survival of only 12 to 15 months ^41^. Diffuse gliomas are among the most difficult cancers to treat, with therapeutic strategies limited to some combination of surgical resection, radiotherapy, or traditional chemotherapy with few if any effective targeted therapies ^42^.

Although the last addition to the treatment repertoire was Temozolomide 17 years ago ^41^, gliomas are among the most deeply genetically characterized of all tumor types thanks to the efforts of multicenter research consortia such as The Cancer Genome Atlas (TCGA) ^43^.

Transcriptomic studies aimed at understanding the pathogenesis of gliomas then take on particular relevance.

Here, we performed an exhaustive transcriptomic study using patient databases (TCGA-LGGGBM^34^) with the aim of characterizing the role of Receptor-interacting serine/threonine protein kinase 1 (RIPK1) in DG pathology.

To carry out our analyses, we classified the samples from DG patients according to the expression levels of RIPK1 and IDH1 status. Firstly, we observed that those patients with high levels of RIPK1 expression had a lower probability of survival compared to those who expressed lower levels of RIPK1. This difference was significant when analyzing all samples together and when analyzing wtIDH samples separately. These results are consistent with previously reported data since RIPK1 is a master regulator of Necroptosis and necroptosis-pathway-associated genes are unfavorable prognostic markers in glioblastoma multiforme ^44, 45^.

Furthermore, we observed significant differences when analyzing RIPK1 expression levels in relation to the histological classification of DG. The same occurred when analyzing the expression of RIPK1 classifying patients according to their IDH1 mutational status. This information together suggests that in more aggressive tumors there is greater expression of RIPK1. Additionally, we observed higher levels of RIPK1 expression in those patients previously treated with radiotherapy, suggesting that RIPK1 could be involved in some mechanism triggered in response to treatment such as inflammation induced by radiotherapy ^46^.

From our differential expression and GO analyses, we selected proliferation, EMT and cell death pathways in order to evaluate the correlation between the expression levels of RIPK1 and the more remarkable genes of these pathways. In this sense, we observed a negative correlation between RIPK1 and most of the genes linked to proliferation, a positive correlation with the key genes involved in EMT and a strong positive correlation between RIPK1 and the genes involved in cell death pathways, especially pyroptosis and necroptosis. These results, taken together with those obtained in relation to prognosis, suggest that the deregulation of some of these pathways could contribute to the lower probabilities of survival observed in those patients with high expression of RIPK1. This is consistent with the results previously reported by Lin et al. ^47^ and Chen et al. ^45^ in which they demonstrate that greater activation of inflammatory cell death pathways correlates with a worse clinical prognosis. At the same time, MPhil et al. ^48^ reported that radiotherapy treatment induces greater activation of pyroptosis and consequently more inflammation, consistent with our findings.

On the other hand, Melo-Lima and collaborators ^49^ used a RIPK1 inhibitor to perform pre-clinical in vitro trials. Treatment with Edelfosine that induces inflammatory cell death combined with the RIPK1 inhibitor produced a redirection of the death pathways, inducing higher levels of apoptosis and lower inflammation.

In relation to the immune response, different authors point out that the IDH mutation in glioma is associated with a reduction in immunological infiltrates leading to a better prognosis in patients with glioma ^50, 51^. In fact Gonzalez et al. ^3^ demonstrated that mIDH gliomas have less immunological infiltration than wtIDH tumors and a greater relative abundance of NK cells ^52, 53^.

Regarding our findings, we found that higher RIPK1 expression correlates with higher immune infiltration in both mIDH and wtIDH tumors. These results could explain the correlation observed between RIPK1 expression and the most aggressive glioma subtypes.

Among the mIDH gliomas, high RIPK1 expression samples exhibited an upregulation of gene signatures of helper and cytotoxic T cells, DCs, macrophages NK cells and neutrophils. On the other hand, high RIPK1 wtIDH gliomas showed an upregulation of macrophages, neutrophils and memory T cells while Tregs and NK cells gene signatures were significantly lower than low RIPK1 tumors. The accumulation of macrophages and neutrophils characterizes the unsolved chronic inflammation involved in the pro-tumorigenic effect of the TIME in glioma ^54^, and thus, it is coherent to find that these gene signatures are upregulated in tumors with such poor prognosis as wtIDH gliomas ^55, 56^.

It is well known that RIPK1 is involved in the inflammatory response and necroptosis^20, 44, 57^. Furthermore, our analysis demonstrated a higher correlation with genes involved in pyroptosis. Several studies have demonstrated that pyroptosis may facilitate the supportive tumor microenvironment ^47, 58^. In this direction, RIPK1 could play a role in sustaining inflammation through the regulation of pro-inflammatory death pathways and collaborate with glioma pathogenesis.

## CONCLUSION

In this study, we proposed an integrative work based in genomic and transcriptomic analysis, with the aim of characterizing the role of the apoptosis inhibitor RIPK1 in Glioma.

All together our *in silico* results suggest that RIPK1 plays an important role in Glioma progression and pathogenesis and could highlight the need to take into account RIPK1 expression levels for decision making to choose the appropriate treatment for each patient or to design novel therapeutic alternatives.

## ACKNOWLEDGEMENTS

We thank the *Agencia Nacional de Promoción Científica y Tecnológica* (ANPCyT), the *Consejo Nacional de Investigaciones Científicas y Técnicas* (CONICET), National University of La Plata (UNLP) and University of Buenos Aires (UBA).

## CONTRIBUTORS

**LCAM:** Conceptualization, Methodology, Writing - original draft. **SMGB:** Methodology, Software, Writing-review & editing. **AM:** Methodology, Validation, Writing -review & editing. **MLS:** Methodology, Writing -review & editing. **NG:** Methodology. **MC:** Conceptualization, Writing -review & editing. **VR:** Supervision, Writing - review & editing. **MLP:** Conceptualization, Project administration, Investigation, Writing - review & editing and Funding acquisition.

## SUPPLEMENTARY MATERIAL

Supplementary material is available online.

